# NIfTI-MRS: A standard format for magnetic resonance spectroscopic data

**DOI:** 10.1101/2021.11.09.467912

**Authors:** William T Clarke, Mark Mikkelsen, Georg Oeltzschner, Tiffany K. Bell, Amirmohammad Shamaei, Brian J. Soher, Uzay E Emir, Martin Wilson

## Abstract

**Purpose:** The use of multiple data formats in the MRS community currently hinders data sharing and integration. NIfTI-MRS is proposed as a standard MR spectroscopy data format, which is implemented as an extension to the neuroimaging informatics technology initiative (NIfTI) format.

Using this standardised format will facilitate data sharing, ease algorithm development, and encourage the integration of MRS analysis with other imaging modalities.

**Methods:** A file format based on the NIfTI header extension framework was designed to incorporate essential spectroscopic metadata and additional encoding dimensions. A detailed description of the specification is provided. An open-source command-line conversion program is implemented to enable conversion of single-voxel and spectroscopic imaging data to NIfTI-MRS. To provide visualisation of data in NIfTI-MRS, a dedicated plugin is implemented for FSLeyes, the FSL image viewer.

**Results:** Alongside online documentation, ten example datasets are provided in the proposed format. In addition, minimal examples of NIfTI-MRS readers have been implemented. The conversion software, *spec2nii*, currently converts fourteen formats to NIfTI-MRS, including DICOM and vendor proprietary formats.

**Conclusion:** The proposed format aims to solve the issue of multiple data formats being used in the MRS community. By providing a single conversion point, it aims to simplify the processing and analysis of MRS data, thereby lowering the barrier to use of MRS. Furthermore, it can serve as the basis for open data sharing, collaboration, and interoperability of analysis programs. It also opens possibility of greater standardisation and harmonisation. By aligning with the dominant format in neuroimaging, NIfTI-MRS enables the use of mature tools present in the imaging community, demonstrated in this work by using a dedicated imaging tool, FSLeyes, as a viewer.

## Introduction

Magnetic resonance spectroscopy (MRS) is a highly flexible technique which can generate a wide range of sensitive and specific imaging contrasts complementary to the typically water-derived contrasts of MR imaging (MRI). MRS allows simultaneous measurement of multiple in vivo metabolite concentrations, which – when combined with additional dynamic contrast encoding techniques – can be used to derive metabolite concentration time courses, metabolite diffusion properties, or in vivo chemical kinetics (1–3).

In-vivo MRS data typically requires a complex data storage array containing multiple dimensions, including spectral frequency (equivalently encoded as a time dimension), spatial encoding, and any dimensions required for ‘dynamic’ encoding. Before pre-processing, additional array dimensions may be required, e.g., to separately store signals from multi-channel receive coils, or multiple transients (4). In addition, any information about the acquisition that is required for interpretation needs to be stored, too, including the transmitter frequency and the signal dwell time, as well as information on the spatial dimensions and orientation of the measurement volume.

To date, there is no established standardised data format for communicating MRS and MR spectroscopic imaging (MRSI) data. Whilst a DICOM standard for MRS(I) exists (5), it is not fully implemented by the vendors. Crucially, DICOM does not provide an intuitive way to store spectroscopic data, nor a standardised method to encode data that requires two or more additional encoding dimensions. As a result, vendors have developed their own separate, proprietary (closed) formats to store raw and processed spectroscopic data. The specific format used is dependent on the scanner software version and local practice. These proprietary formats vary greatly in the degree in which data has been subjected to inline-processing, i.e. whether data has been spatially reconstructed, which (and how) metadata is stored, and in the way the storage is formatted.

The lack of a standard data format hinders the use of MRS in several ways:

1. **It impedes integration with other imaging modalities**. Without standardised encoding of spatial orientation and position, registration with other modalities requires a per-data-format solution. This hinders both co-analysis of spectroscopic data with other modalities, and leveraging of other modalities in reconstruction and processing of the typically low-signal MRS.
2. **It impedes the consistent analysis of data**. Any spectroscopic analysis program must implement dedicated interpretation modules for each format. Development of new modules is time consuming and relies on expert knowledge of a format. This results in incomplete coverage of the formats by any one analysis pipeline. Thus consistent (and comparable) analysis is prevented.
3. **It impedes the creation of new, discrete, specialised tools**. Without a standard storage format, processing and analysis of spectroscopic data often occurs in a single monolithic process, frequently relying on local analysis pipelines and depending on the local MRS expert to create them. This impedes modular processing, which is important for development and uptake of new acquisition and analysis methods, as well as improving existing ones. The barrier to creating new tools is high if implementation of all processing steps is required. Furthermore, quantifying improvements from modifying a discrete step is difficult if it is inseparable from other steps.
4. **It raises the difficulty of sharing data**. Compared to other modalities, sharing MRS data is not straightforward. With the diversity of storage formats, especially across platforms, users cannot reliably read and interpret data received from other users. Data sharing repositories are required to handle mixed data types, which are processed to varying levels and have varying metadata. This creates enormous sharing friction, and discourages researchers from sharing. Anonymisation tools required for ethical public sharing of data require per-format implementation.

Combined, these factors significantly raise the barrier to adoption and use of MRS, especially for non-expert users. Compared to other MRI modalities, MRS analysis workflows remain highly customised and specific, and require unique MRS expertise on site.

To address these issues, we propose a single data format based on the Neuroimaging Informatics Technology Initiative (NIfTI) format (6) for storing single-voxel, spectroscopic imaging, and unlocalised spectral data. We call the proposed format NIfTI-MRS. The NIfTI file format is the standard format for storing anatomical, functional, diffusion and quantitive MRI and arterial spin labelling data in the MR neuroimaging research domain. NIfTI has provided a cornerstone for analysis across neuroimaging, allowing integrated analysis of modalities.

The proposed format will:

- provide a simplified pathway from scanner to final analysis,
- enable interoperability and modularity of analysis programs,
- enable easier display and co-interpretation with other modalities, and
- establish a format for data sharing.

In addition, the proposed format is designed to provide a simple anonymisation procedure and flexible storage of meta-data, further removing friction from the process of sharing.

To facilitate adoption, this work describes the implementation of an open-source command-line conversion program capable of converting many original formats to the proposed format. The program, *spec2nii*, provides single-point conversion for all spatially reconstructed data (including single-voxel data) from 14 different formats alongside anonymisation scripts and manual editing tools for NIfTI-MRS.

By aligning MRS with the most widely used neuroimaging format, NIfTI-MRS will also allow researchers to create comprehensive MRS research designs that incorporate different modalities with ease. To demonstrate this, we have created an FSLeyes (FMRIB software library, University of Oxford)(7) plugin to visualise multi-dimensional NIfTI-MRS data alongside structural MRI and results from MRS fitting.

## Methods

### The NIfTI-MRS data format

A brief description of the proposed standard (version 0.3) is included here. The full standard is provided as supporting information, and the latest version can be found at Reference (8).

#### Design

The NIfTI-MRS format is designed to contain single-voxel, contiguous multi-voxel, and unlocalised (or partially localised) time-domain MRS data in up to three spatial dimensions (I.e. MRSI encoded data). Optionally, NIfTI-MRS can encode up to three additional data dimensions, e.g. for arrays of interrelated signals. The standard is designed with low minimum conformance metadata requirements to simplify adoption, while providing for more complicated metadata requirements in the full format.

The NIfTI format contains three sections: the data header, optional header extensions, and the data block. The proposed NIfTI-MRS format comprises a NIfTI-2-formatted file with a mandatory header extension formatted according to the JavaScript Object Notation (JSON) standard (9,10).

In NIfTI-MRS, the NIfTI data header (11,12) structure is identical to the one used for structural or functional MRI data, although some values are constrained or re-utilised for spectroscopy-specific purposes. For example, the dwell-time of the time domain data is stored in *pixdim[4]* (with units are specified in the *xyzt_units* field). Spatial position encoding, i.e. dimensions and orientation of the measurement volume, is implemented as in the NIfTI specification, and a default value of *pixdim* is specified for unlocalised data. Finally, only complex datatypes may be specified in the *datatype* field (e.g. “DT_COMPLEX”, 32). The NIfTI-MRS standard is versioned, and the version is specified in the *intent_name* field (in the format *mrs_vMajor_minor*).

The NIfTI-MRS data block is used to store up to seven-dimensional complex time-domain data. The first four dimensions are required for a valid NIfTI-MRS file: the first three dimensions are used for spatial encoding (*x, y*, and *z* coordinates), the fourth dimension is used to store the time-domain FID (or echo). All three spatial dimensions have a size of one for single-voxel data. The remaining three dimensions (fifth, sixth, and seventh) are optional. They flexibly encode different dynamic aspects, with the specific purpose and interpretation of each dimension being documented in the header extension.

The header extension has the official NIfTI identification code *44*, “*NIFTI_ECODE_MRS*”. It comprises key-value pairs formatted according to the JSON standard, which can be arbitrarily nested, if necessary. The header extension contains the minimum necessary metadata required for meaningful interpretation of the spectroscopic data, additional information about the optional higher encoding dimensions, and further MRS-specific metadata.

#### Header Extension Metadata

The header extension contains four types of metadata key:

1. The two <u>mandatory</u> keys *SpectrometerFrequency & ResonantNucleus*. These must be present in all NIfTI-MRS formatted files, because they are a necessary requirement for correct reconstruction and interpretation of the time-domain data.
2. Higher encoding dimension information. Three keys are defined per additional encoding dimension *(n=5,6,7): dim_{n}, dim_{n}_info, and dim_{n}_header*. The first, *dim_{n}, is mandatory if the dimension is used. dim_{n} takes the value of a list of predefined tags (e*.*g. DIM_COIL or DIM_EDIT)* to identify the purpose of the dimension. A *DIM_USER_{0-2}* tag allows for user-specified purposes, which can be described using the *dim_{n}_info field*. The *dim_{n}_header* tag enables each element of a dimension to be associated with different values of metadata keys.
3. Standard-defined metadata. These keys correspond to well-defined and frequently used sequence, hardware, or subject data. They are defined in the NIfTI-MRS standard and may not be redefined. These keys are optional.
4. nUser-defined metadata. Keys can be arbitrarily defined by users to store unusual metadata, that not foreseen in the standard, or that for which no fixed format or recommendations exist. User-defined JSON metadata permits additional fields for user-defined keys, encouraging in-place documentation to aid interpretation of the keys. These keys are entirely optional.

#### Spatial information

Orientation, position, and voxel size information are stored in the NIfTI-MRS header in accordance with the NIfTI standard. Conformance is achieved in the header either when:

1. *qform_code* is set > 0,
2. the second to fourth elements of *pixdim* are set to the appropriate voxel dimensions, or to a default of 10 m,
3. *quatern_b, quatern_c, quatern_d, qoffset_x, qoffset_y, qoffset_z* are set,
4. a valid value of *qfac* is set.

or:

1. *qform_code* is set = 0 (*NIfTI_XFORM_UNKNOWN*),
2. the second to fourth elements of *pixdim* are set to the appropriate voxel dimensions, or to a default of 10 m.

The former *pixdim* option is suitable for data which has a meaningful spatial position, and the latter for data with no real-world position, e.g. simulated data. Use of the default pixdim size (10 m) indicates a dimension has no localisation, or or that it has poorly defined extent (e.g. localisation provided only by the limited extent of coil sensitivity).

The NIfTI format in general (and therefore also NIfTI-MRS) cannot store spatially non-contiguous data (i.e. data with a gap between voxels or slices) in a single file. Distinct contiguous volumes need to be stored separately.

#### Anonymisation of protected health information

To ease the process of anonymisation, all standard-defined metadata keys are marked as privacy-sensitive or not privacy-sensitive in the standard. User-defined metadata may be self-marked as privacy-sensitive by appending *‘private_’* to the start of the key or any nested key within the definition.

Anonymisation of data stored as NIfTI-MRS is simplified through two features:

1. Only a subset of metadata is retained in the conversion to NIfTI-MRS. This metadata is selected and only incorporates that which is well defined.
2. Anonymisation tools acting on NIfTI-MRS can reliably identify sensitive fields which have been converted using their definition in the standard.

#### Processing provenance

Pre-processing steps applied to the data can be optionally recorded in the header extension. The type of pre-processing applied, the program used, the program version, and any additional information provided by the pre-processing algorithm can be stored sequentially in the ‘ProcessingApplied’ field. Provenance is not provided by the NIfTI-MRS data standard itself, but either requires adequate implementation in relavent software packages or manual addition by users.

#### Phase & frequency conventions

The NIfTI-MRS standard defines a strict frequency and phase convention for data. This convention follows the conventions of Levitt (13). In this convention the absolute frequency scale should increase from left to right. For nuclei with a gyromagnetic ratio greater than zero this corresponds to resonances from nuclei with less shielding (more deshielding), which therefore experience a higher magnetic field, appearing on the left. I.e. they have more negative (higher magnitude) Larmor frequencies, noting w = -yB_O_. This results in a typically displayed chemical shift (‘ppm’) axis increasing from right to left. A description of this convention in the spectral time-domain is provided in the specification.

### Software implementation

To promote the use of NIfTI-MRS the standard has been implemented into software for conversion, visualisation, and data input-output. Functions for loading, writing, and visualizing NIfTI-MRS data have been created for common programming languages and integrated into open-source analysis pacakages.

#### Conversion to NIfTI-MRS

An open-source conversion program *spec2nii* has been created. *Spec2nii* reads vendor-proprietary, DICOM, and processing toolbox formats and generates NIfTI-MRS-formatted files. The program can also inspect, edit and anonymise existing NIfTI-MRS files. The program operates on the command-line, is written in Python, and developed as a public, open-source resource.

#### Visualisation using FSLeyes

To enable visualisation of the multi-dimension NIfTI-MRS format, an FSLeyes plugin has been created. FSLeyes is the free, open-source FMRIB Software Library (FSL) image viewer. The developed plugin extends FSLeyes to interpret the NIfTI-MRS format and enables display of single and multi-voxel spectroscopy stored in the format.

## Results

### The NIfTI-MRS data format

The specification for the NIfTI-MRS data format is available with this document as supporting information, and online at Reference (8). To assist in the interpretation of the standard, both online explanatory documentation (https://wtclarke.github.io/mrs_nifti_standard/) and example data (14) have been created. Figure 2 shows extracts of NIfTI-MRS header extensions from four of the ten example datasets. These four examples demonstrate: the structure of the header extension, the flagging of privacy-sensitive data, dynamic header values describing different conditions in one of the additional data dimensions (here spectral editing), and records of processing provenance.

**Figure 1:**
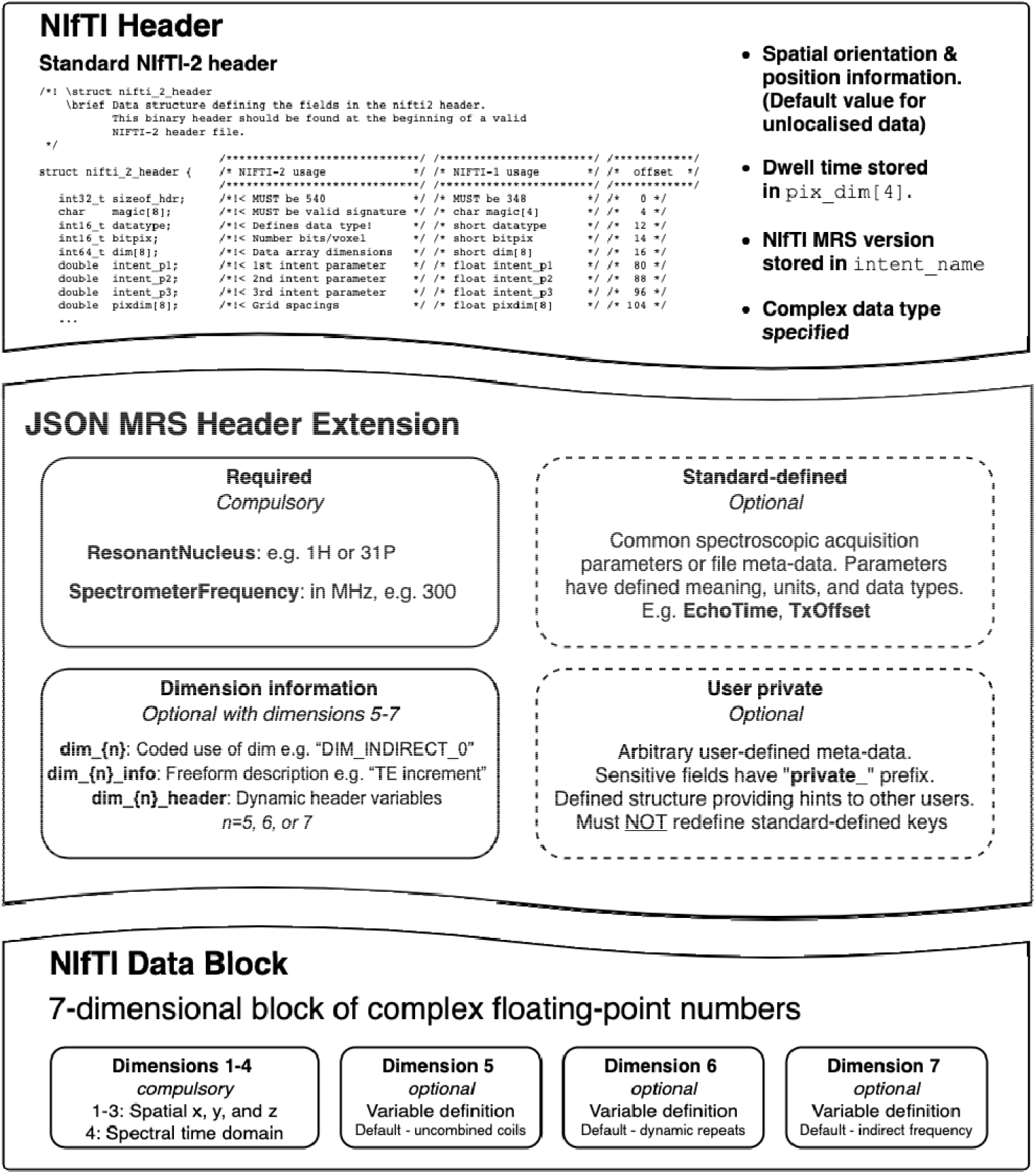
Schematic representation of the proposed NIfTI-MRS format. The format utilises the native NIfTI header and Data Block, whilst using a JSON-formatted NIfTI header extension to store additional required metadata.

**Figure 2:**
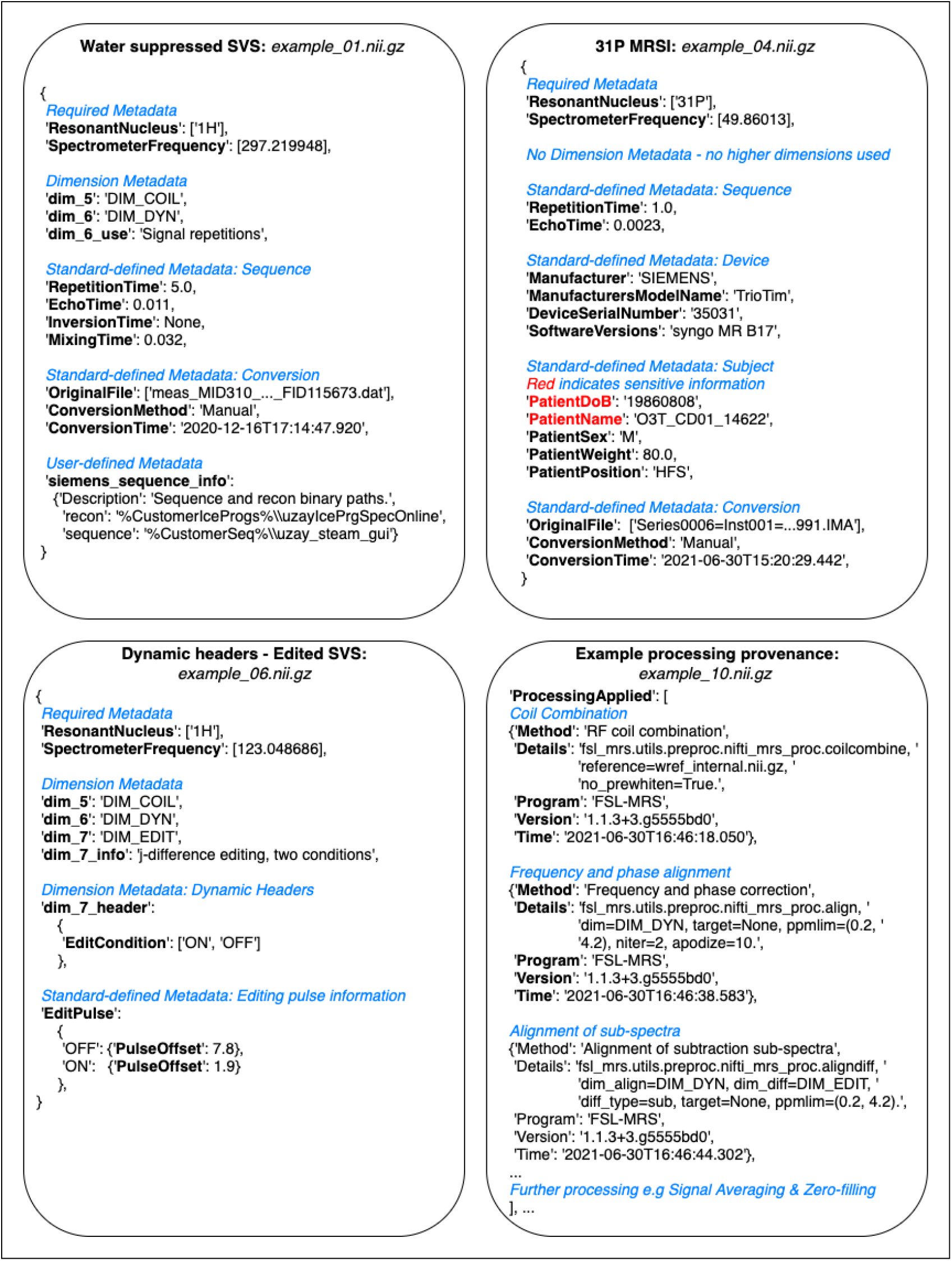
Extracts of NIfTI-MRS JSON-formatted header extensions for four different pieces of example data. The full example data is available from reference (14). Each example demonstrates a different aspect of the header extension format. Figure annotations are shown as blue italicised text. **A:** Structure of a header extension of^1^ H single-voxel data before pre-processing. **B:** Header extension for processed^31^ P MRSI, including fields that are marked for anonymisation (red). **C:** Example of dynamic header fields indicating an editing condition stored in the 7^th^ dimension. **D:** Extract of the processing provenance in a MEGA-PRESS (15) sequence pre-processed using FSL-MRS (16).

### Software implementation

#### Conversion to NIfTI-MRS

The conversion program *spec2nii* has been created and released as a Python package. The package is developed online (https://github.com/wtclarke/spec2nii), and is available from the PyPI and Conda package managers. Spec2nii provides automatic or semi-automatic conversion of 14 data formats (vendor-proprietary and DICOM) to NIfTI-MRS (Table 1).

**Table 1:** List of spec2nii supported formats in version 0.3.4. WIP = Work in progress

| Vendor / software | Format | File extension | Spectroscopy formats handled |  |  | Automatic Orientation | Notes |
| --- | --- | --- | --- | --- | --- | --- | --- |
|  |  |  | SVS | MRSI | FID |  |  |
| Siemens | Twix | .dat | ✓ | × | ✓ | ✓ | VB&VE baselines |
| Siemens | DICOM | .ima | ✓ | ✓ | ✓ | ✓ |  |
| Philips | SPAR/SDAT | .SPAR/.SDAT | ✓ | × | ✓ | ✓ |  |
| Philips | data-list | .data/.list | ✓ | × | ✓ | ✓ |  |
| Philips | DICOM | .dcm | ✓ | × | ✓ | WIP |  |
| GE | pfile | .7 | ✓ | ✓ | ✓ | ✓ | Per-sequence mapping required |
| UIH | DICOM | .dcm | ✓ | ✓ | ✓ | ✓ |  |
| Bruker | 2dseq | - | ✓ | ✓ | ✓ | ✓ | PV 5.1, 6.0, 6.0.1, 7.0.0, 360 |
| Bruker | fid | - | ✓ | × | ✓ | WIP | PV 5.1, 6.0, 6.0.1, 7.0.0, 360 |
| Varian | fid | - | ✓ | × | ✓ | WIP |  |
| LCModel | RAW/H2O | .RAW/.H2O | ✓ | × | ✓ | × |  |
| jMRUI | Text | .txt | ✓ | × | ✓ | × |  |
| jMRUI | MRUI | .mrui | ✓ | × | ✓ | × |  |
| - | ASCII | .txt | ✓ | × | ✓ | × |  |

Figure 3 outlines the proposed use cases of *spec2nii* and NIfTI-MRS enabled by this work:

1. varied input data is passed to spec2nii,
2. spec2nii identifies the appropriate reconstruction and conversion pathway,
3. conversion is carried out, and a NIfTI-MRS file is returned,
4. pre-processing is carried out using NIfTI-MRS as an intermediary format,
5. pre-processed data is stored or further analysed (fitted).

**Figure 3:**
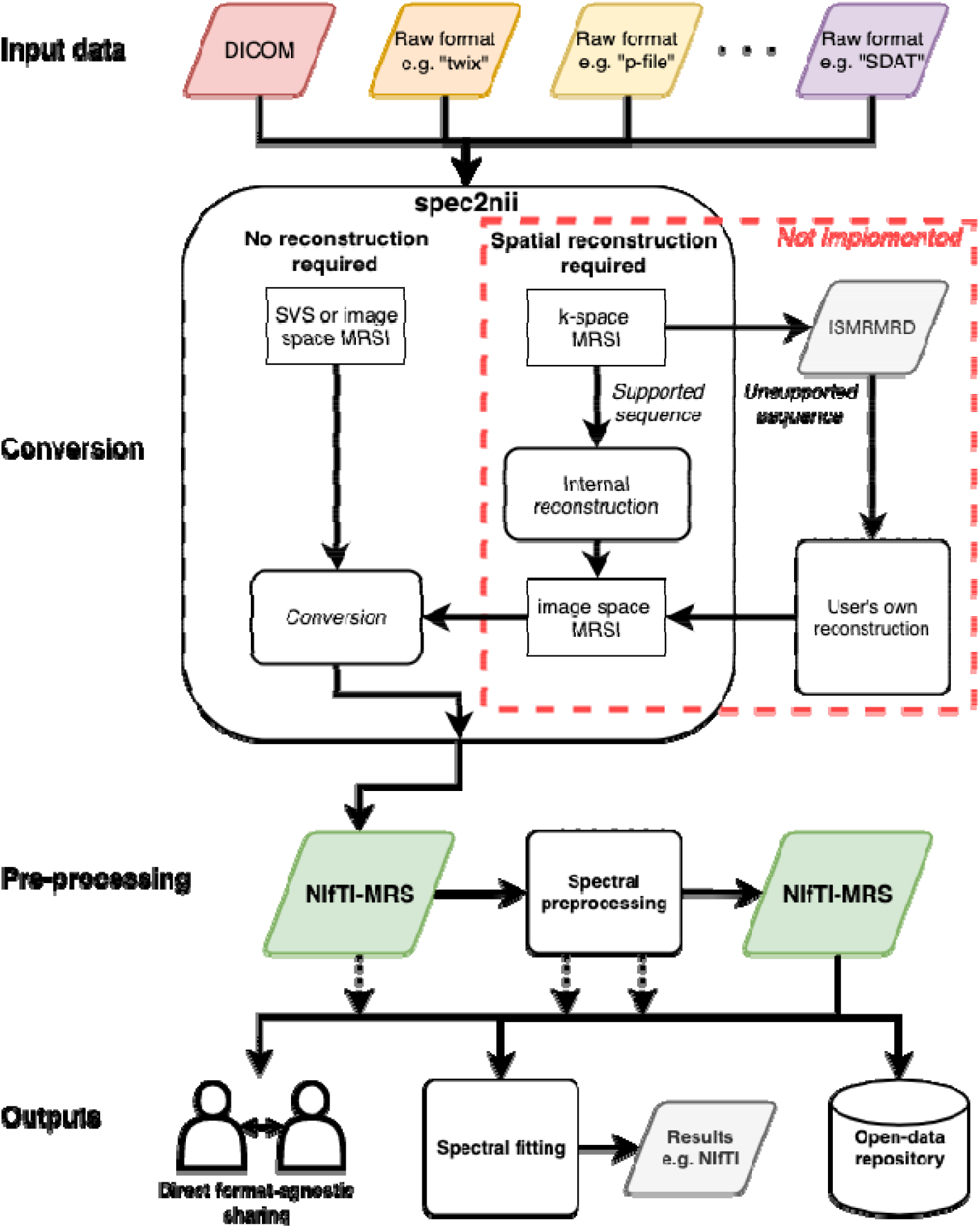
Proposed MRS and MRSI processing pipelines using NIfTI-MRS and incorporating conversion in spec2nii. In the proposed pipeline, raw data from a variety of formats (e.g. DICOM, Siemens “Twix .dat”, GE “p-file” or Philips “SDAT/SPAR”) are converted to NIfTI-MRS using spec2nii. Subsequently pre-processing can be applied, with both the input and results stored in NIfTI-MRS. Data can be shared with other users or a data repository in a format-agnostic way at any stage of the pre-processing pipeline. The pre-processed NIfTI-MRS file can then be passed on to modelling software. spec2nii can convert unlocalised, single-voxel, and spatially reconstructed MRSI. In the future spec2nii will also convert MRSI stored in k-space for certain common supported sequences. For other sequences (e.g. those with non-cartesian trajectories) a pipeline incorporating the ISMRMRD format (17) and third-party reconstruction is proposed. The red box indicates software yet to be implemented.

The data at any stage after conversion (#3) can be shared; whether that is the unprocessed converted data, fully pre-processed, or partially processed data.

For this work only the conversion of spatially reconstructed data has been implemented in *spec2nii*. Data that are stored in a k-space representation cannot currently be converted. In the future some spatially unreconstructed data from standard vendor-supplied MRSI sequences will be handled by *spec2nii*. For data requiring specialist reconstruction, e.g. those with non-cartesian trajectories, we propose a future pathway incorporating conversion to ISMRMRD format (17) and third-party reconstruction provided by the sequence developers.

The formats supported by spec2nii are summarised in Table 1. *Spec2nii* carries out automatic spatial orientation calculations for seven of the fourteen supported formats.

#### NIfTI-MRS I/O and Compatibility

At Reference (18) minimal examples of NIfTI-MRS file readers have been provided in four common programming languages (Java, Matlab, Python, and R). The examples exploit the availability of robust NIfTI I/O libraries in those programming languages.

Support for NIfTI-MRS has been established in five open-source spectroscopy analysis packages: FSL-MRS (16), Osprey (19), Spant (20), Vespa (21) [https://github.com/vespa-mrs/vespa], and FID-A (22). Each package has input-output compatibility with the standard. Example short-TE STEAM and water reference data at 7T, provided in the NIfTI-MRS format and processed in FSL-MRS, spant and Osprey (with Osprey using FID-A as foundation), are shown in Figure 4. The spatial location of the single voxel is shown overlaid on a NIfTI structural image in each package. This demonstrates the benefit for interoperability of different software solutions, and the potential to form the foundation of a mutually compatible, multi-language analysis software ecosystem

**Figure 4:**
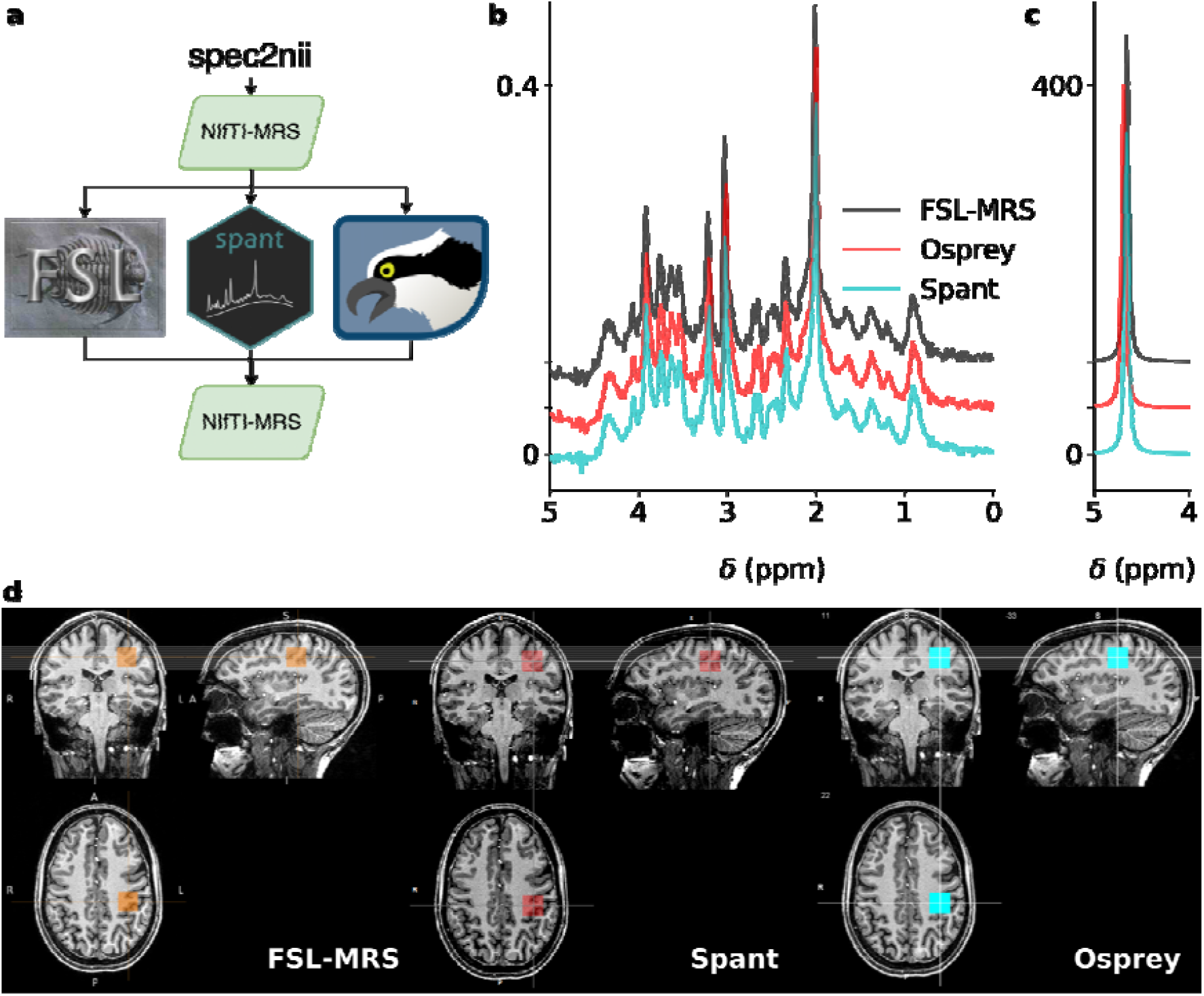
NIfTI-MRS single voxel spectroscopy data processed in three spectroscopy toolboxes supporting input and output of NIfTI-MRS data. Raw data converted to NIfTI-MRS format by spec2nii was loaded and pre-processed in each of the toolboxes, before being written back out to NIfTI-MRS. Comparison of water-suppressed (b) and water-reference (c) data processed in each toolbox is simple using the pipeline in a. The output NIfTI-MRS data is easy to present alongside structural data stored in NIfTI format using a variety of MRS analysis software (d).

#### Visualisation in FSLeyes

Visualisation of NIfTI-MRS-formatted data is possible using FSLeyes and the NIfTI-MRS plugin for FSLeyes. The plugin implements:

- a pan-able & zoomable display for spectra,
- display of NIfTI-MRS headers,
- automatic calculation of the chemical shift axis,
- display of individual spectra stored in the higher dimensions (5^th^-7^th^ dimensions),
- interactive zeroth and first-order phasing of spectra, and
- easy comparison of spectra from different voxels.

The plugin is maintained at https://git.fmrib.ox.ac.uk/wclarke/fsleyes-plugin-mrs and is available from the Pypi and Conda package managers. It operates using the FSLeyes interface, is written in python, is open source, and available under the BSD 3-Clause licence. FSLeyes is available under the Apache License, Version 2.0 (7).

This work enables NIfTI-MRS formatted data to be displayed alongside arbitrary MRI and other modality imaging data formatted as NIfTI. It also enables the visualisation of multidimensional NIfTI-MRS data. Figure 5 shows two examples of NIfTI-MRS data displayed in FSLeyes using the plugin.

**Figure 5:**
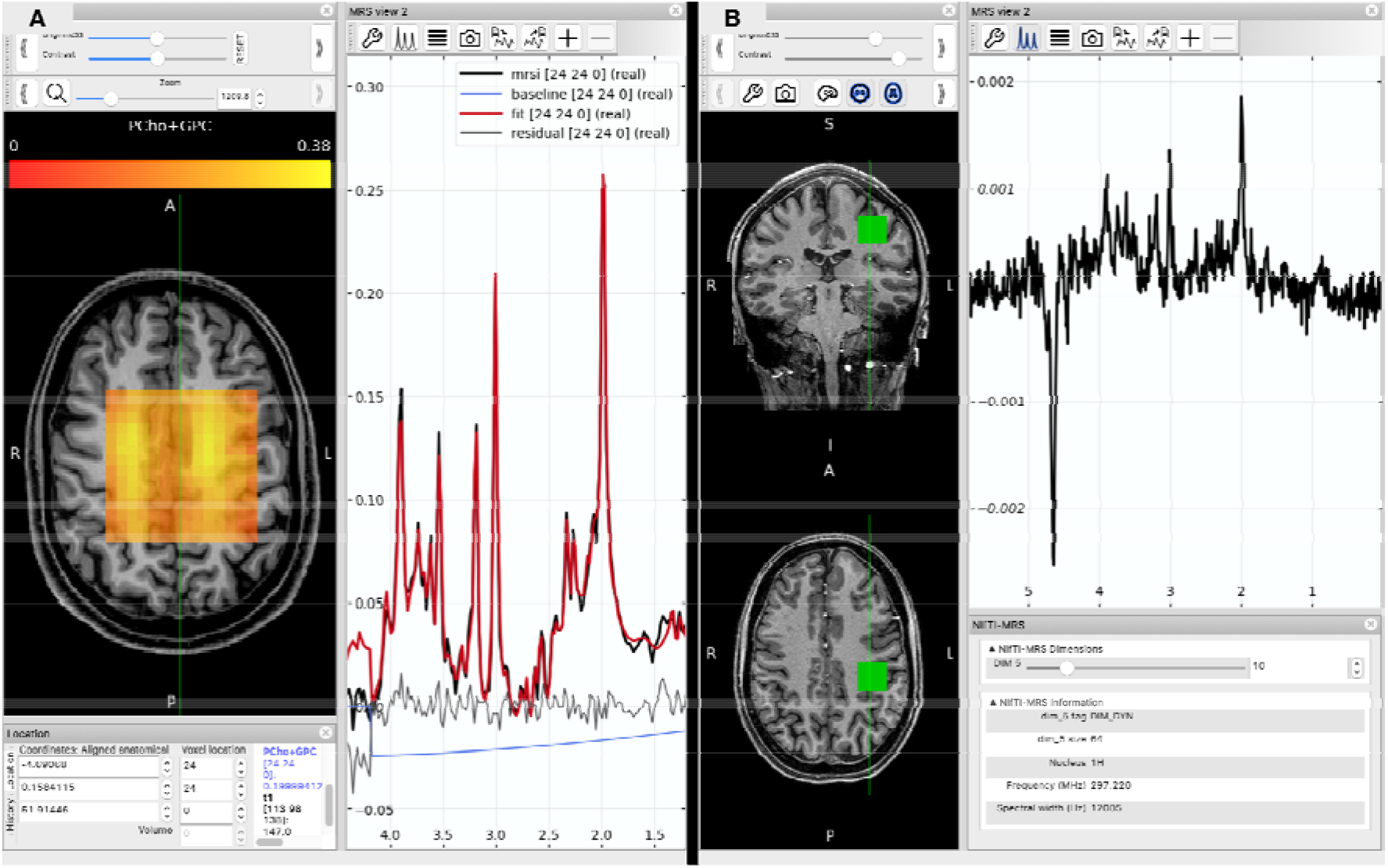
A: MRSI data displayed in FSLeyes using the NIfTI-MRS plugin. In addition to the NIfTI-MRS data, results from spectral fitting (also stored in NIfTI format) are displayed overlaid on the spectrum, to the left a metabolite map of total choline is displayed overlaid on a T1w structural image. The spectrum display automatically displays voxel-wise spectrum and fits as the cursor is moved over the orthographic display. B: Partially pre-processed single voxel data displayed in FSLeyes alongside corresponding structural data. Under the spectral view a dedicated NIfTI-MRS panel displays selected information from the header extension and allows the user to select elements from the higher (5^th^ -7^th^) dimensions. Here the 10^th^ (of 64) temporal average stored in dimension 5 is displayed.

## Discussion & Conclusion

In this work we have proposed NIfTI-MRS as a new standard data storage format for MRS and MRSI. The standard was initially developed and agreed by a group of MRS physicists and researchers with experience in MRS analysis software use and development. Following a draft specification, subsequently feedback was sought from MRS software developers, experts in the field, and the wider technical MRS community. The standard and associated files are maintained online at Reference (18).

NIfTI-MRS can provide a simplified analysis pathway, provide a path for interoperability of analysis programs, and greatly simplifies display and co-interpretation of MRS data alongside other modalities. In addition, the standard fulfils several other desirable objectives:

- A standard data storage format is a prerequisite for incorporating MRS data into standardised data structuring schemes, such as BIDS (23).
- Similarly, a standard format allows for extensive and easy data archiving and sharing in open science databases, such as OpenNEURO (24,25) or MRS-specific databases(26). One study (27) has already utilised NIfTI-MRS for the purpose of releasing study data on XNAT central (https://central.xnat.org/data/projects/PN21). And another has recently submitted data formatted as NIfTI-MRS to the National Institute of Mental Health Data Archive (28).
- Standard header definitions and processing provenance make it easier for users to comply with minimum reporting standards consensus statements (29), and to reproduce and recreate data analysis workflows simply by extracting the provenance information from a NIfTI-MRS file they received

The developers of NIfTI-MRS have sought to make the standard as accessible as possible. *Spec2nii* implements a mature open-source program suitable for both one-off and batch conversion tasks. NIfTI I/O libraries are available in all common programming languages, and minimal examples of readers have been provided. Support for NIfTI-MRS is already available in five analysis packages (with further packages implementing the standard currently). This represents a critical mass of actively developed processing tools.

To date *spec2nii* handles 14 different formats, however complete coverage of sequences across such a diverse range of formats cannot be guaranteed. Currently data must be in a spatially reconstructed format before conversion. Development of *spec2nii* has relied on a community-led model, with users contributing examples, test data, and code contributions to the developers.

In this work FSLeyes has been extended with a plugin to create an NIfTI-MRS compatible data viewer. The authors consider this just an example of the existing tools present in the MR imaging community that can be easily leveraged for MRS and MRSI by using NIfTI-MRS. NIfTI-MRS mitigates the difficulty of aligning and registering MRSI data with MRI (and other modality) data, easing the simultaneous use of imaging and spectroscopic data to further methodological and analysis techniques in both fields.

The NIfTI-MRS standard is versioned, enabling backwards-compatible future extension of the standard. The authors propose that the ISMRM MRS Study Group Committee for Code and Data Sharing (https://www.mrshub.org), established in 2020 as a permanent standing committee with rotating members, maintain oversight of the standard and any future development.

Establishing a standardised data format for MRS is one requirement for MRS to be used for large scale cross-centre, or population, studies. The NIfTI format provided this for neuroimaging studies such as the Human Connectome Project (30). Furthermore, the uptake of the NIfTI-MRS standard shows the possibility of establishing an ecosystem of open-source analysis toolboxes similar to that which has benefitted functional neuroimaging [e.g. AFNI, FSL, SPM] (31–33). Nonetheless, this work does not tackle other key aspects such as communication of MRS fitting results.

NIfTI-MRS has been designed to both promote the use of MRS in biomedical research, but to also ease the technical development of MRS analysis in the wider field of biomedical imaging. The standard and associated tools are developed in an open-research context. Further use and development of NIfTI-MRS and associated tools are dependent on the expertise and contributions of the community, including MR hardware vendors. To-date the community has enthusiastically done so, enabling the rapid development of the standard and its inclusion in multiple software tools.

## Supporting information

Specification

## Acknowledgements

The authors thank all contributors of test data and source code to the *spec2nii* program, especially Jack Miller and Tomáš Pš orn. Also, Paul McCarthy, the FSLeyes developer, for his extensive advice in creating the NIfTI-MRS visualisation plugin.

William T Clarke is supported by funding from the Wellcome Trust and the Royal Society (102584/Z/13/Z). The Wellcome Centre for Integrative Neuroimaging is supported by core funding from the Wellcome Trust (203139/Z/16/Z).

Georg Oeltzschner receives support from the National Institute of Health (K99/R00 AG062230, S10 OD021648, P41 EB031771, P41 EB015909, R01 EB016089, R01 EB023963, and R01 EB028259).

Amir M Shamaei has received funding from the European Union’s Horizon 2020 research and innovation program under the Marie Sklodowska-Curie grant agreement No 813120.

## Supporting information

1. nifti_mrs_specification_0_5_1.pdf – Specification of the NIfTI-MRS format

