## Supplementary material for "NIfTI-MRS: A standard format for magnetic resonance spectroscopic data": Specification

### NIfTI MRS format specification

Version 0.5

#### 1 Purpose

MR vendors export MRS data in a number of different formats, each of which are liable to change due to regular scanner software updates. Supporting a variety of (often undocumented) formats across a variety of MRS techniques (SVS, MRSI, fMRS, edited MRS…) places a significant and unnecessary burden on MRS software developers and researchers. Furthermore it makes effective data sharing extremely difficult. The DICOM format has been successful in enabling interoperability between MR scanners and hospital PACS, allowing MR images from all vendors to be seamlessly transferred across networks for viewing and archiving. Whilst a DICOM standard for MRS exists, it is not fully implemented by the vendors, and other DICOM 'hacks' for MRS have resulted in crucial information being stored inconsistently between vendors or contained in private tags (which require expertise to interpret). A more complete implementation of the DICOM MRS standard remains an important goal to enable routine use of MRS in a clinical radiology department, however the format is poorly suited for research use due to its clinical focus and unnecessarily complex features - such as nested sequences. Crucially, DICOM MRS does not provide an intuitive or standardised way to store data which has two or more dynamically changing parameters, for example individual coil elements or dynamic scans, both of which are important in MRS research.

Recently a number of open-source MRS analysis packages have been developed, presenting a timely opportunity to create a standard format for storing MRS data with an emphasis on ease of implementation and support for the full range of MRS acquisition types. The existing NIfTI format has been chosen to store MRS data due to its:

1. simplicity;
2. excellent availability of supporting libraries in Python, MATLAB, R, JAVA, and C;
3. ubiquity in the neuroimaging community; and
4. pre-existing support by the major neuroimaging software packages (FSL, SPM, AFNI).

The Neuroimaging Informatics Technology Initiative (NIfTI) file format was originally developed in 2003 to address the needs of MR neuroimaging researchers, and has since become the standard format for storing anatomical, fMRI, diffusion MRI, qMRI and ASL data in this domain. Whilst the NIfTI format was primarily developed for neuroimaging and fMRI, the format is sufficiently flexible to store a wide range of imaging data - including MRS.

In this document we propose a standard for the storage of MRS data as a NIfTI file with the following objectives:

1. Provide a standard entry point for researchers using MRS analysis software.
2. Improve interoperability between MRS software packages.
3. Reduce duplication of efforts by MRS software developers to support the numerous existing formats.
4. Provide support for the full range of MRS acquisition types.
5. Provide a standard format for sharing MRS data.
6. Align MRS data storage practices more closely to other (neuro)imaging techniques to improve compatibility with existing platforms and initiatives such as https://openneuro.org/ and https://bids.neuroimaging.io/.

#### 2 Specification

The NIfTI format defines a binary file with filename extension ".nii" which optionally may be compressed e.g. ".nii.gz". A NIfTI file comprises a header (Official definition of the nifti1 header / nifti2.h) followed by a multidimensional array of numerical data points. Optionally, one or more 'header extensions' may follow the header. The format of a header extension is indicated by a predefined 'ecode' number. NIfTI header keys will be distinguished in this document using a highlighted font. For example, the first header value corresponds to the size of the header: `int sizeof_hdr`. All indexes are counted from zero, i.e. `pix_dim[0]` is the first element of the array and {N} should be substituted as 0, 1, 2….

The NIfTI MRS format inherits all definitions from the NIfTI standard, with the following additional requirements:

1. The intent name (`char intent_name[16]`) specifies conformance to a version of this standard using the format "mrs_vM_m" (M=major version, m=minor version). For example "mrs_v0_2".

2. Data points must be stored as 64bit (or higher precision) complex as indicated by `int datatype`. I.e. DT_COMPLEX, DT_COMPLEX64, or DT_COMPLEX128.
3. NIfTI header orientation fields must be complete and populated as described in §2.2 Spatial Positioning and Orientation.
4. Required MRS metadata key/value pairs must be stored in a NIfTI header extension as a JavaScript Object Notation (JSON) formatted string. See *§2.3 MRS metadata*.

This standard prefers the NIfTI-2 format (https://nifti.nimh.nih.gov/nifti-2). Compared to NIfTI-1, NIfTI-2 implements a simplified header capable of storing 64-bit values. The NIfTI-1 format is acceptable for this standard but should be avoided where possible.

#### 2.1 Complex time series data

MRS data in this standard is stored as complex time domain data. Data points must be stored as 64bit (or higher precision) complex as indicated by `int datatype`. I.e. DT_COMPLEX, DT_COMPLEX64, or DT_COMPLEX128. The dwell time (time domain increment) is stored as the fifth element of the `pixdim`array. The units of this dimension are specified by the fourth to sixth bits of the `xyzt_units` field, and should be set to one of seconds, milliseconds or microseconds (NIfTI_UNITS_SEC, NIfTI_UNITS_MSEC, NIfTI_UNITS_USEC).

Phase and frequency conventions for the storage of the complex data are given in Appendix A.

#### 2.2 Spatial positioning and orientation

To conform with this standard, orientation, position, and voxel size information must be stored in the NIfTI header with no deviation from the NIfTI standard. I.e. conformance is achieved when in the NIfTI header either:

1. `qform_code` is set > 0,
2. the second to fourth elements of `pixdim`are set to the appropriate voxel dimensions or to a default of 10 m (10000 mm) for unlocalised dimensions`,`
3. `quatern_b, quatern_c, quatern_d, qoffset_x, qoffset_y, qoffset_z`are set;
4. A valid value (1 or -1) of `qfac` is set. `qfac`is stored as the first element of`pixdim.`

or:

1. `qform_code` is set = 0 (NIfTI_XFORM_UNKNOWN),
2. the second to fourth elements of `pixdim`are set to the appropriate voxel dimensions or to a default of 10 m (10000 mm) for unlocalised dimensions.

The former option is suitable for data which has a meaningful spatial position, and the latter for data with no real-world position, e.g. simulated data. Use of the default `pixdim` size (10 m) indicates that the data collected has no or poorly-defined (e.g. coil only) localisation in that dimension.

The NIfTI format contains an additional way of specifying the spatial information. An affine matrix may be specified directly in the `srow_x, srow_y` and `srow_z` fields. This standard supports the original intended use of the sform matrix and `sform_code`field to permit orientation for the data to an additional coordinate system e.g. a defined "standard" space such as MNI 152.

The spatial units of the second, third and fourth elements of `pixdim`and the `qoffset` fields are specified by the first three bits of the `xyzt_units` field which should be set appropriately to meters, millimeters or micrometers (`NIfTI_UNITS_METER, NIfTI_UNITS_MM, NIfTI_UNITS_MICRON`).

##### 2.2.1 Spatially non-contiguous data

The NIfTI standard cannot store spatially non-contiguous data. Data with slice-gaps or non-parallel slices cannot be encoded using the "qform" or "sform" methods independently. Data of this type should be stored in multiple files separated into contiguous spatial regions.

#### 2.3 MRS metadata

In addition to the standard header of the binary NIfTI file, other MRS specific information can be stored in a NIfTI header extension. This metadata must be stored as key-value pairs in a JSON formatted text string. The header extension concept is specified in the official NIfTI documentation. The JSON formatted extension contains a set of key-value pairs (in any order), with some key names following a fixed dictionary defined herein.

MRS data may be stored for a variety of purposes, from only requiring access to the raw FID data points, to storing geometry and acquisition details necessary to perform metabolite simulation and partial volume correction of metabolite concentrations.[1, 2] This standard defines a minimum level of conformance required for meaningful interpretation of the data. Subsequently the standard defines a series of MRS metadata keys and groups of keys which are required to carry out certain analysis operations. Users may also include their own metadata keys, if not already a part of this standard, but must not redefine those defined in this standard.

The header extension consists of two mandatory fields (`int esize, ecode`) stored in the first 8 bytes, followed by the UTF-8 encoded byte representation of the JSON formatted metadata string. The whole extension (mandatory fields + JSON) must be padded to an integer multiple of 16 bytes. The size is then set as the value of `esize.` I.e.`esize` must be a positive integer multiple of 16. The `ecode` for the MRS NIfTI specification header extension is 44 ("NIfTI_ECODE_MRS").

THe JSON "null" value is permitted within the meta-data, though all keys apart from those in §2.3.1 are optional. JSON arrays should not be of mixed types.

###### 2.3.1 Required MRS metadata

For conformance to this standard the JSONheader extension must contain the following required fields:

| Key | Datatype | Description |
|---|---|---|
| SpectrometerFrequency | Array of numbers | Precession frequency in MHz of the nucleus being addressed for each spectral axis. See DICOM tag (0018,9098). |
| ResonantNucleus | Array of strings | See DICOM tag (0018,9100). Must be one of the DICOM recognised nuclei "1H", "3HE", "7LI", "13C", "19F", "23NA", "31P", "129XE" or one named in the specified format. I.e. Mass number followed by the chemical symbol in uppercase. |

More than one value is permitted for each of these keys to allow for multi-nuclei experiments. For example, a 1H-13C HSQC acquisition at 7T would have the following JSON entries:

```
"SpectrometerFrequency":[300, 75.5]
"ResonantNucleus":["1H", "13C"]
```

JSON metadata specified as an array of values (e.g. "Array of numbers") must be given as an array even for single element arrays. I.e. "SpectrometerFrequency":[300] not "SpectrometerFrequency":300 in the previous example.

The other essential MRS parameter, dwell time, is stored in `pixdim[4]`.

###### 2.3.2 Data dimensionality

It is possible to store up to 7 dimensions of MRS data points in a NIfTI compatible file - with the first 4 dimensions being mandatory. The first three dimensions index the X, Y and Z spatial coordinates of the voxel (each dimension having a length of 1 for SVS) with the fourth dimension storing the complex time-domain data. For example, a reconstructed 2D MRSI scan acquired with a 16x16 matrix in the X-Y plane and 1024 complex data points in the time-domain would have the following dimensions 16x16x1x1024. Note that NIfTI MRS data should be stored in the image domain (rather than as spatial frequencies) and the fourth dimension must be in the time-domain (rather than the frequency-domain).

`short dim[8]` contains the sizes of the data array dimensions, with the first element (`dim[0]`) specifying the number of dimensions. In the previous example `dim[0]` would be set to 4 - which is the smallest permissible value for NIfTI MRS data. Data may be optionally stored across the remaining 3 dimensions, with these being used to store the following information by default:

- `dim[5]` - data from individual receiver coil elements,
- `dim[6]` - dynamic scans, more than one FID acquired sequentially (either prior to averaging, or for fMRS),
- `dim[7]` - the indirect dimension - required for storing MRS with two frequency axes (2D NMR).

For example SVS data collected from a 32ch head coil, with 1024 complex data points and 128 repetitions would have the following dimensions 1x1x1x1024x32x128 with `dim[0]` having a value of 6. Data dimensions of 1x1x1x1024x32x128x1 with `dim[0]` having a value of 7 would be equally valid.

The optional data dimensions may also be used to store alternative information (or deviate from the default ordering) by specifying values for the following keys in the JSON extension header: "dim_5", "dim_6" and "dim_7". These values have a data type of string and {N} denotes a zero-based index:

| Value | Meaning |
|---|---|
| "DIM_COIL" | For storage of data from each individual receiver coil element. |
| "DIM_DYN" | For storage of each individual acquisition transient. E.g. for post-acquisition B0 drift correction. |
| "DIM_INDIRECT_{N}" | The indirect detection dimension - necessary for 2D (and greater) MRS acquisitions. |
| "DIM_PHASE_CYCLE" | Used for the time-proportional phase incrementation method. |
| "DIM_EDIT" | Used for edited MRS techniques such as MEGA or HERMES. |
| "DIM_MEAS" | Used to indicate multiple repeats of the full sequence contained within the same original data file. |
| "DIM_USER_{N}" | User defined dimension. |
| "DIM_ISIS" | Dimension for storing image-selected in vivo spectroscopy (ISIS) acquisitions. |

For example a 2D MRS scan with a FID length of 1024 points and 64 increments in the indirect dimension could be stored with data dimensions 1x1x1x1024x64, provided the following was specified in the JSON sidecar:

```
"dim_5": "DIM_INDIRECT_0"
```

The use of each dimension as specified by keys in the "dim_5", "dim_6" and "dim_7" fields, can be optionally made more explicit by utilising the optional "dim_5_info", "dim_6_info" and "dim_7_info" fields. These fields may be freeform strings:

```
"dim_5": "DIM_INDIRECT_0"
"dim_5_info": "Echo time increment"
```

If the dimension is used to increment over defined metadata key(s) then this can be conferred by using the optional "dim_5_header", "dim_6_header" and "dim_7_header" fields with values stored as JSON arrays:

```
"dim_5": "DIM_INDIRECT_0"
"dim_5_info": "Echo time increment"
"dim_5_header": {"EchoTime": [0.03, 0.04, …, 0.10] }
```

More than one metadata key can be specified using a JSON object:

```
"dim_5_header": {"EchoTime":  [0.03, 0.04, …, 0.10],
                 "RepetitionTime":  [1.0, 1.10, …, 2.00]}
```

For more information and examples see §2.3.5 *Dynamically changing metadata*. Both the dim_{N}*info and dim*{N}_header keys are optional. Note that the sixth to eighth elements of pixdim (`pixdim[5], pixdim[6], pixdim[7]`) are not used for this purpose.

###### 2.3.3 Standard defined metadata

In addition to the metadata required for minimum conformance and interpretability (§2.3.1), further metadata might be needed for meaningful interpretation of the stored data. For example storing information on basis simulation; relaxation correction; subject, sequence and scanner information; or editing pulse information.

Therefore this standard defines a set of optional metadata key-value pairs which encode common acquisition and sequence parameters. In this specification we explicitly define the names, data types and descriptions of a number of these keys (see §5 Appendix B). Users may define their own keys in addition to those specified here, but must not change the meaning of existing keys in Appendix B.

**Anonymisation of data**

Standard-defined metadata keys are given a standard anonymisation flag which marks the key for deletion or retention upon anonymisation. This information is given in Appendix B.

##### 2.3.4 User-defined metadata

As described in §2.3.3 users may define their own metadata for inclusion in the JSON formatted MRS header extension. These user defined keys cannot redefine those contained in Appendix B, those defined in §2.3.1 *Required MRS metadata*, or the NIfTI header.

To aid understanding of user-defined metadata these parameters should be grouped by purpose into a JSON object. Within the JSON object a reserved "Description" key allows for a free-form text description of the other key-value pairs in the object.

At either the top level or within the json object the key prefix "private_" indicates that the key should be removed upon anonymisation.

| Key | Datatype | Description |
|---|---|---|
| {user_defined_key} | Any, including nested JSON structures. | Arbitrary user defined meta-data. |
| private_{user_defined_key} | Any, including nested JSON structures. | Arbitrary user defined meta-data. Should be removed upon anonymisation. |
| Description | string | Freeform description of the user defined key. |

Single-valued user meta-data should also use this format, providing a description to identify the purpose of the user-defined field. For example:

```
"Excitation pulse duration":
    { "Value" : 3.0,
      "Description" : "Duration of the excitation pulse."}
```

Or for multiple related user-defined fields:

```
"Excitation pulse information":
    { "Duration": 3.0,
      "Pulse name": "SINC",
      "Description" : "Excitation pulse information. Duration in ms."}
```

##### 2.3.5 Dynamically changing metadata

The optional 5th, 6th and 7th data dimensions can store data acquired under dynamically changing acquisition conditions, e.g. with spectral editing or non-uniformly varying parameters. For example, the 5th dimension could be tagged DIM_INDIRECT_{N} or DIM_USER{N}, specifying a description in dim_5_info, and associating one or more metadata keys by specifying dim_5_header.

The values of the associated metadata for each dimension index can then be specified in two ways.

1. For standard-defined metadata, the value at each dimension index can be described:

    1. fully, by using a JSON array of length equal to the size of the dimension,
    2. in abbreviated form, using a JSON object with fields "start" and "increment".

    The units and form of each element should match the metadata key specification in appendix B. Option b is only suitable for meta-data taking point numeric values at fixed increments.

2. For user-specified metadata the format of §2.3.4 should be followed, using a JSON object, including a "Description" field with the iterable array or short format contained in "Value".

Examples

Example data is included in the official standard git repository alongside the descriptions in this document.

*Example 1: j-difference editing utilising a user-defined meta-data structure to define the position of the editing pulse. The data contains two conditions "ON" and "OFF" contained in dim 7 after an uncombined coils dimension and a dimension containing repeated averages.*

```
"dim_5": "DIM_COIL"
"dim_6": "DIM_DYN"
"dim_7": "DIM_EDIT"
"dim_7_info": "j-difference editing, two conditions"
"dim_7_header": {"EditCondition": [ "ON", "OFF"]},
```

*Example 2: j-evolution utilising a uniformly incremented TE evolution. The Echo time is specified in the shortened object form in dimension 6 (after a coils dimension).*

```
"dim_5": "DIM_COIL"
"dim_6": "DIM_INDIRECT_0"
"dim_6_info": "Incremented echo time for j-evolution"
"dim_6_header": {"EchoTime": {"start": 0.03, "increment": 0.01}}
```

*Example 3: j-evolution utilising a non-uniform incremented TE evolution. The Echo time is specified fully as an array in dimension 6 (after a coils dimension).*

```
"dim_5": "DIM_COIL"
"dim_6": "DIM_INDIRECT_0"
"dim_6_info": "Incremented echo time for j-evolution"
"dim_6_header": {"EchoTime": [0.035, 0.036, 0.037, 0.04, … 0.125]}
```

*Example 4: A fingerprinting based example. Dimension 5 is used to index changes across three standard-defined metadata keys (EchoTime, RepetitionTime, and ExcitationFlipAngle) as well as a user defined inversion/phase condition.*

```
"dim_5": "DIM_USER_0"
"dim_5_info": "Acquisition index with variable TE, TR, flip-angle and pulse offset."
"dim_5_header":
    {"EchoTime": [0.000, 0.001, 0.002, 0.005, … 0.090],
     "RepetitionTime": [0.0, 0.1, 0.2, 0.1, … 0.0],
     "ExcitationFlipAngle": [10, 20, 30, … 100],
     "Inv_condition":
        {"Value" : [0, 180, …, 180],
         "Description" : "User defined inversion condition."}}
```

#### 3 Resources

###### 2.5.1 NIfTI

https://brainder.org/2012/09/23/the-nifti-file-format/

https://nipy.org/nibabel/nifti_images.html#the-nifti-header

https://www.nitrc.org/docman/view.php/26/204/TheNIfTI1Format2004.pdf

https://nifti.nimh.nih.gov/pub/dist/src/niftilib/nifti1.h

###### 2.5.2 BIDS

https://bids-specification.readthedocs.io/en/stable/

https://www.nature.com/articles/sdata201644

###### 2.5.3 DICOM

http://dicom.nema.org/medical/dicom/current/output/chtml/part03/sect_C.8.14.html

###### 2.5.4 JSON

http://json.org

### 4 Appendix A: Complex data phase conventions

NIfTI-MRS Follows the conventions of Levitt (The Signs of Frequencies and Phases in NMR. Journal of Magnetic Resonance 1997;126:164–182 doi: 10.1006/jmre.1997.1161). In this convention the absolute frequency scale should increase from left to right, noting that $\omega$=-$\gamma B_0$.

This means for nuclei with a gyromagnetic ratio > 0 (e.g. $^{1}$H) this corresponds to resonances from nuclei with less shielding (more deshielding), which therefore experiencing a higher magnetic field, appearing on the left. I.e. they have more negative (higher magnitude) Larmor frequencies.

We also define the DFT as

$$A_k = \sum_{m=0}^{n-1} a_m \exp\left\{-2\pi i \frac{mk}{n}\right\} \qquad k = 0, \ldots, n-1.$$

to match the definition of Levitt , Numpy and MATLAB.

*Note: In literature the frequency axis is often also plotted with a different frequency scale showing more positive relative frequencies on the left. This preserves the orientation of the spectrum but corresponds to a scale equal to* $|\omega|-|\omega_{ref}|$.

To comply with this convention we define a right-handed coordinate system.

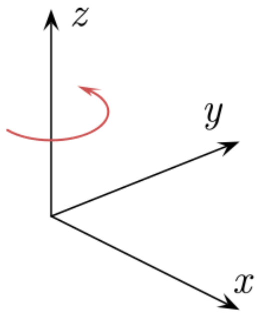


with x corresponding to real values, y imaginary values, and z time. In this frame a positive rotation is defined as counter-clockwise looking down the time axis (shown in red). The spectral data points are ordered in sequence of increasing time.

In this coordinate frame:

- data arising from nuclei with a positive gyromagnetic ratio should be stored so that positive relative frequencies (on the absolute frequency scale and relative to the spectrometer reference frequency) appear to have a positive (counter-clockwise) rotation.
- data arising from nuclei with a negative gyromagnetic ratio should be stored so that positive relative frequencies (on the absolute frequency scale and relative to the spectrometer reference frequency) appear to have a negative (clockwise) rotation.

### 5 Appendix B: Specified JSON metadata tags

This appendix contains the standard-defined metadata keys. Keys have been grouped here for legibility but are not nested within the header extension.

#### 5.1 MRS specific tags

| Key | Datatype | Unit | Description |
|---|---|---|---|
| EchoTime | number | s | Time from centroid of excitation to start of FID or centre of echo. Units: Seconds |
| RepetitionTime | number | s | Sequence repetition time. Units: Seconds |

| | | | |
|---|---|---|---|
| InversionTime | number | s | Inversion time. Units: Seconds |
| MixingTime | number | s | Mixing time in e.g. STEAM sequence. Units: Seconds |
| AcqusitionStartTime | number | s | Time, relative to EchoTime, that the acquisition starts. Positive values indicate a time after the EchoTime, negative indicate before the EchoTime, a value of zero indicates no offset. Units: Seconds |
| ExcitationFlipAngle | number | degrees | Nominal excitation pulse flip-angle |
| TxOffset | number | ppm | Transmit chemical shift offset from SpectrometerFrequency |
| VOI | Array of numbers | - | VoI localisation volume for MRSI sequences. Stored as a 4 x 4 affine using identical conventions to the xform NIfTI affine matrix. Not defined for data stored with a single spatial voxel. |
| WaterSuppressed | boolean | - | Boolean value indicating whether data was collected with (True) or without (False) water suppression. |
| WaterSuppressionType | string | - | Type of water suppression used. |
| SequenceTriggered | boolean | - | Boolean value indicating whether the sequence is triggered. If triggered the repetition time might not be constant. |

##### 5.2 Scanner information

| Key | Datatype | Description | Anon (Y/N) |
|---|---|---|---|
| Manufacturer | string | Manufacturer of the device. DICOM (0008,0070). | N |
| ManufacturersModelName | string | Manufacturer's model name of the device. DICOM (0008,1090). | N |
| DeviceSerialNumber | string | Manufacturer's serial number of the device. DICOM (0018,1000). | Y |
| SoftwareVersions | string | Manufacturer's designation of the software version. DICOM (0018,1020) | N |
| InstitutionName | string | Institution's Name. DICOM (0008,0080). | Y |
| InstitutionAddress | string | Institution's address. DICOM (0008,0081). | Y |
| TxCoil | string | Name of transmit RF coil. | N |
| RxCoil | string | Name of receive RF coil | N |

##### 5.3 Sequence information

| Key | Datatype | Description | Anon (Y/N) |
|---|---|---|---|
| SequenceName | string | User defined name. DICOM (0018,0024). | N |
| ProtocolName | string | User-defined description of the conditions under which the Series was performed. DICOM (0018,1030). | N |

##### 5.4 Subject information

| Key | Datatype | Description | Anon (Y/N) |
|---|---|---|---|
| PatientPosition | string | Patient position descriptor relative to the equipment. DICOM (0018,5100). Must be one of the DICOM defined code strings e.g. HFS, HFP. | N |
| PatientName | string | Patient's full name. DICOM (0010,0010). | Y |
| PatientID | string | Patient's ID. DICOM (0010,0020). | Y |

| | | | |
|---|---|---|---|
| PatientWeight | number | Weight of the Patient in kilograms. DICOM (0010,1030). | N |
| PatientDoB | string | Date of birth of the named Patient. YYYYMMDD. DICOM (0010,0030). | Y |
| PatientSex | string | Sex of the named Patient. 'M', 'F', 'O'. DICOM (0010,0040) | N |

##### 5.5 Data provenance and conversion metadata

| Key | Datatype | Description | Anon (Y/N) |
|---|---|---|---|
| ConversionMethod | string | Program used for conversion. May include additional information like software version. | N |
| ConversionTime | string | Time and date of conversion. ISO 8601 compliant format. I.e. "YYYY-MM-DDThh:mm:ss.sss" | N |
| OriginalFile | Array of strings | Name and extension of the original file(s) | Y |

##### 5.6 Spatial information

| Key | Datatype | Description | Anon (Y/N) |
|---|---|---|---|
| kSpace | Array of booleans | Three element list, corresponding to the first three spatial dimensions. If True the data is stored as a dense k-space representation. | N |

##### 5.7 Editing Pulse information structure

A formalism for containing information about editing conditions.

| Key | Datatype | Description | Anon (Y/N) |
|---|---|---|---|
| EditCondition | Array of strings | List of strings that index the entries of the EditPulse structure that are used in this data acquisition. Typically used in dynamic headers (dim_N_header). | N |
| EditPulse | JSON Object | Structure defining editing pulse parameters for each condition. Each condition must be assigned a key. | N |

Each condition is listed as an element in an array of JSON objects under the key "EditPulse". Each value of the object is defined with the following optional fields.

| Key | Datatype | Description |
|---|---|---|
| PulseOffset | number | ppm |
| PulseAmplitude | Array of numbers | Amplitude modulation in Hz |
| PulsePhase | Array of numbers | Phase modulation in radians |
| PulseDuration | number | Pulse duration in seconds |
| Nucleus | string | Nucleus |

For example with MEGA-PRESS data the header extension could contain the following:

```
"dim_7": "DIM_EDIT"
"dim_7_info": "j-difference editing, two conditions"
"dim_7_header": { "EditCondition" : [ "ON", "OFF"]},
```

```
"EditPulse": { "ON" : { "PulseOffset" : 1.9},
               "OFF" : { "PulseOffset" : 7.8}}
```

This would identify the last (7th) dimension as having a size of two, with the first index corresponding to the on-resonance saturation condition, and the second index to the control condition.

##### 5.8 Processing Provenance

| Key | Datatype | Description | Anon (Y/N) |
|---|---|---|---|
| ProcessingApplied | Array of JSON Objects | Array of objects as defined below. Describes and records the processing steps applied to the data. | Y |

An array for storing a record of processing steps applied to the data. Each element of the array takes the structure described below. Processing steps can be specified in an arbitrary separate file indicated by the 'Link' field.

| Key | Datatype | Description |
|---|---|---|
| Time | string | Time and date of processing step. ISO 8601 compliant format. I.e. "YYYY-MM-DDThh:mm:ss.sss". |
| Program | string | Processing program used |
| Version | string | Version of program used |
| Method | string | Description of processing step applied. See list in this section for fixed definitions. |
| Details | string | Optional program specific details. |
| Link | string | File path (either/or) |

The 'Method' field can take arbitrary string values but where possible users should adhere to the keywords given in tables 2,3 and 4 of Near *et al* (NMR in Biomedicine e4257 doi:10.1002/nbm.4257). These keywords are reproduced below.

| | |
|---|---|
| Eddy current correction | Signal averaging |
| Frequency and phase correction | Phasing |
| Alignment of subtraction sub-spectra | Apodization |
| Nuisance peak removal | Zero-filling |
| RF coil combination | Subtraction / Addition of sub-spectra |

#### Licence

This specification is released under the Creative Commons Attribution 4.0 International (CC BY 4.0) Licence.

#### Notes

1: Gasparovic C, Song T, Devier D, et al. Use of tissue water as a concentration reference for proton spectroscopic imaging. Magnetic Resonance in Medicine 2006;55:1219–1226 doi:10.1002/mrm.20901.

2: Gasparovic C, Chen H, Mullins PG. Errors in 1H-MRS estimates of brain metabolite concentrations caused by failing to take into account tissue-specific signal relaxation. NMR in Biomedicine 2018;31:e3914 doi:10.1002/nbm.3914.
